# Congruence between noise and plasticity in protein expression

**DOI:** 10.1101/2024.08.18.608377

**Authors:** Saburo Tsuru, Chikara Furusawa

## Abstract

Gene expression responds to various perturbations, such as mutations, environmental changes, and stochastic perturbations. The variability in gene expression levels differs among genes, influencing the availability of adaptive variants or mutants and thereby affecting nongenetic and genetic adaptations. Different types of variability are interdependent, suggesting global canalization/decanalization against different perturbations and a common underlying mechanism. Despite this, the relationship between plasticity (variability in response to environmental changes) and noise (variability among cells under the same conditions) in gene expression remains debatable. Previous studies reported a positive correlation between plasticity and noise, but these variabilities are often measured at different levels: plasticity at the mRNA level and noise at the protein level. This methodological discrepancy complicates the understanding of their relationship. We investigated this by measuring protein expression levels of essential and nonessential genes in *Escherichia coli*. Using flow cytometry, we quantified noise and plasticity from the same dataset. Essential genes exhibited lower noise and plasticity than nonessential genes. Nonessential genes showed a positive correlation between noise and plasticity, while essential genes did not. This study provides empirical evidence of essentiality-dependent coupling between noise and plasticity in protein expression, highlighting the organization of different types of variabilities.

## Introduction

Gene expression changes in response to various perturbations, such as genetic mutations and environmental shifts. Even among genetically identical cells with identical environmental histories, gene expression can vary, a phenomenon known as stochastic noise in gene expression^1^. The variability in gene expression levels in response to these perturbations differs among genes^2–5^. For example, mutations can cause expression levels of some genes to vary more frequently than others, even when mutations are randomly distributed^3,4^. This gene-to-gene difference in variability is not only a consequence of past evolution but can also affect the availability of new adaptive mutants, thereby influencing the rate and direction of de novo phenotypic evolution^4,6–8^. These evolutionary impacts are not limited to variability in response to mutations; environmentally induced phenotypic variability, or plasticity, has the potential to initiate and direct adaptive evolution^6–9^ through processes such as genetic assimilation^10^. Even stochastic noise can facilitate evolution^11^, such as seen in phenomena like partial penetrance^12^. Therefore, understanding the organization of variability in the gene expression of individual genes is of great importance in evolutionary biology.

Different types of variability are closely related and often interdependent^6–8,13–16^. Environmentally induced transcriptional variation often mirrors genetically induced variation in two key aspects^4,7,8^. First, genes that are sensitive in expression levels to environmental perturbations also tend to be sensitive to genetic perturbations^3,4,17–19^. For example, genes essential for cell division and maintenance tend to exhibit lower transcriptional variability in response to both environmental and genetic perturbations in *E. coli* and yeast. In contrast, genes involved in amino acid catabolism, a crucial metabolic pathway for coping with fluctuations in nutrient conditions, tend to exhibit higher transcriptional variability in response to different types of perturbations in *E. coli*. Secondly, a consistent directionality in covariation of expression levels among genes is shared by environmental and genetic perturbations^5^. In *E. coli*, for example, a bunch of genes involved in starvation responses tends to covary in the same direction in response to both environmental changes and mutations^4,20^. These two types of similarities imply global canalization/decanalization^9,13,14^ and suggest a common mechanism underlying the bias in phenotypic variability against different types of perturbations^4,13,21^. Therefore, without a thorough understanding of the relationships among different types of variability, our comprehension of complex adaptations influenced by various perturbations remains incomplete^6^.

Despite the implication of global canalization suggested by the similarity in transcriptional variability between environmental and genetic perturbations, the relationship between plasticity and noise in gene expression levels remains debatable^3,22–24^. Although several studies have reported a positive correlation between plasticity and noise in gene expression levels^3,4,23,25^, these two types of variability are often measured using different methods: plasticity at the mRNA level, termed transcriptional plasticity, and noise at the protein level, termed protein noise. This methodological discrepancy complicates the exploration of the molecular mechanisms and the evolutionary origins of the relationship between noise and plasticity. Interestingly, previous studies have demonstrated that the coupling between protein noise and transcriptional plasticity is less applicable to essential genes^23,24^, suggesting that congruence between noise and plasticity in expression levels is detrimental for essential genes and that the relationship between these two types of variabilities is an evolvable trait^24^. However, protein expression levels are not determined solely by mRNA expression level but also by multiple factors, such as mRNA structure^26^ and codon usages^27^. Additionally, the ratio of protein expression levels to mRNA expression levels varies among genes^28^. These complexities make it difficult to predict the essentiality-dependent correlation at the protein level. Therefore, quantifying noise and plasticity at the same gene products is crucial for understanding their relationship, global canalization, and underlying molecular mechanisms.

To address this issue, we investigated the relationship between noise and plasticity in protein expression levels. We used a subset of yellow fluorescent protein (YFP) fusion strains of *Escherichia coli*^28^ to measure protein expression levels of essential and nonessential genes. Using flow cytometry, we quantified the mean and cell-to-cell heterogeneity (noise) in protein expression across different nutrient conditions. These conditions were designed to perturb both expression levels and growth rates greatly according to a previous study^4,29^. Crucially, we quantified both noise and plasticity from the same dataset to ensure the reliability of the relationship between noise and plasticity. We found that noise, plasticity, and their relationship depend on gene essentiality. Essential genes tended to exhibit lower noise and plasticity than nonessential genes. For nonessential genes, there was a positive correlation between noise and plasticity, whereas essential genes showed no significant correlation. This study provides the first direct empirical evidences of essentiality-dependent coupling between noise and plasticity in protein expression levels, highlighting organization of different types of variabilities.

## Results

### Measurement of protein expression levels at the single-cell level across different nutrient conditions

We utilized a library of yellow fluorescent protein (YFP) fusion strains of *E. coli*^28^ to explore protein expression levels (**Fig. 1a**). Each chromosomal copy of genes was labeled with a YFP fusion. To ensure reliable measurements of YFP fluorescence using flow cytometry, we selected strains exhibiting moderate fluorescence levels based on the previous data^28^. Additionally, we filtered out stains exhibiting localized fluorescent foci within the cytosol to avoid potential aggregation of YFP-fused proteins, which could interfere with the accurate measurement of YFP fluorescence. Gene essentiality was based on a previous study^30^, which identified 248 genes designated as essential for cellular growth across different datasets. As a result, we randomly selected 89 strains (67 nonessential and 22 essential genes, respectively, **Supplementary Table S1**) and cultured them under different nutrient conditions. These conditions included three carbon sources (glucose, glycerol, and mannose) and eight supplement groups (casamino acids, cocktails of amino acids with different combinations, nucleotides, vitamins, and trace elements) (**Fig. 1b**). Using combinations of these supplement groups, we constructed nine supplement conditions for each carbon source (**Supplementary Table S2**), distinguished by environmental ID. For instance, ID 01 comprised trace elements, vitamins, nucleotides, and casamino acids. We defined supplement richness based on the number of supplement groups, ranging from 1 to 8, where a higher number represents greater supplement richness. These nutrient conditions were designed to obtain different growth rates, established in a previous study^29^. We confirmed that the growth rates ranged from 0.26 to 0.93 (**Fig. 1c**). We note that two environmental conditions (environmental ID=09,10) were identical in supplement richness (five), but differed in combinations of amino acids (6A and 6B) while maintaining the same number of amino acids (six).

**Fig. 1:**
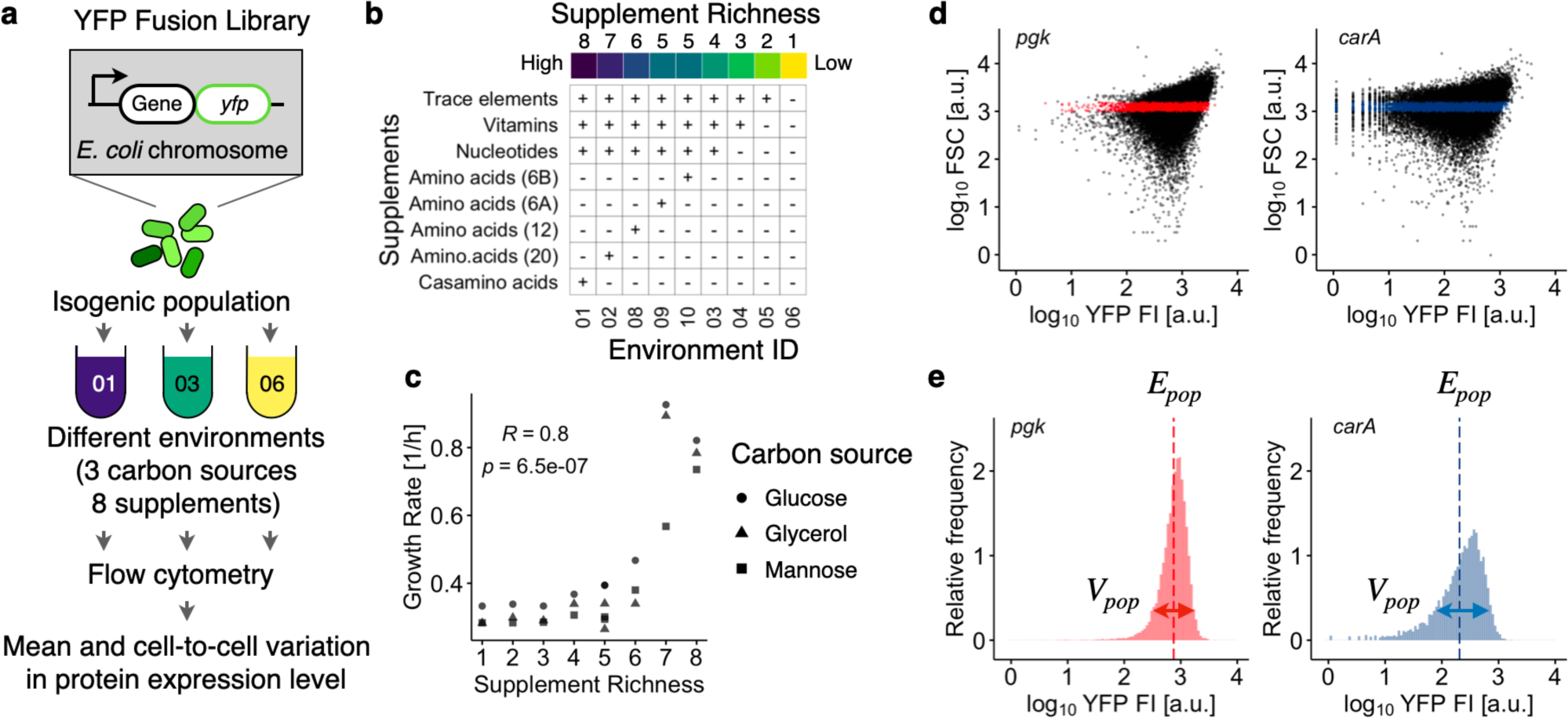
Schematics of experimental workflow. **a**, Schematic diagram illustrating the profiling YFP expression levels at the single-cell level. A total of 89 strains, including 22 essential and 67 nonessential genes, from the YFP fusion library of *E. coli* were analyzed. Each strain carries a YFP-fused ORF at its natural chromosomal location. Strains were cultured under various nutrient conditions, combining three carbon sources with eight supplements, and analyzed using flow cytometry during the exponential growth phase. **b**, Combinations of eight supplements used in nutrient conditions. Supplement richness was defined by the number of supplements. The plus (+) and minus (−) signs indicate the presence or absence of the corresponding supplements (rows) in the nutrient conditions (columns). **c**, Relationship between cell growth rate and supplemental richness. The inset represents Spearman’s rank correlation coefficients (R) and p-value. **d**–**e**, Results for two representative strains carrying YFP-fused pgk (an essential gene, left) and carA (a nonessential gene, right) cultured under a common condition (Environment ID: 10; carbon source: glucose). **d**, Relationship between forward scattering light (FSC) and YFP fluorescence intensity (YFP FI) of individual cells (dots). Cells within the common narrow FSC gate are colored red or blue. The YFP FI after background fluorescence subtraction is shown. Strains with fewer than 4,000 cells within the gates were excluded from subsequent analysis. **e**, YFP distributions of the gated cells in panel **d**. The means (dashed vertical lines) and standard deviations, labeled as E_pop_ and V_pop_, respectively, were calculated on the logarithmic scale. The scale of horizontal arrows representing V_pop_ is exaggerated for visualization.

We measured protein expression levels for each strain during the exponential growth phase across different nutrient conditions using flow cytometry. The YFP fluorescence intensity (YFP FI) reflects the total fluorescence emitted from a single cell, with larger cells generally exhibiting higher YFP FI. To compensate for this cell-size dependency, we filtered the cells using a narrow gate of forward scattering (FSC) (**Fig. 1d**). The gate was defined for each nutrient condition because bacterial cell size depends on nutrient conditions^29^. Clonal cell populations exhibit huge cell-to-cell heterogeneity in gene expression even under identical environmental conditions, resulting in long-tailed or log-normal-shaped distributions in protein numbers^28,31^. This stochastic nature often results in rare cells exhibiting very highly expression levels within cell populations, which greatly influences the variance or standard deviation of protein number distributions. To ensure robust measurement of variation in fluorescent distributions, the YFP FI, after subtracting the autofluorescence intensity, was log_10_-transformed and used to calculate the means (E_pop_) and standard deviations (V_pop_) for each population (**Fig. 1e**), following a conventional method employed in previous studies^25,32^ using flow cytometry for the analysis of protein noise. The E_pop_ and V_pop_ values of each strain in each nutrient condition represent the clonal population mean and cell-to-cell heterogeneity in protein expression levels of the labeled gene, respectively. We averaged E_pop_ and V_pop_ among biological replicates. Using these statistics, we quantified plasticity and noise in protein expression levels as described in the following subsections.

### Plasticity in protein expression levels

We first examined the plasticity in protein expression levels for each gene across different nutrient conditions. To assess the impact of nutritional perturbations on changes in protein expression, we focused on the population means of protein expression levels, E_pop_s, across the various nutrient conditions. A principal component analysis (PCA) revealed that the major axis (PC1) accounted for 66% of the total variation of E_pop_ across these conditions (**Fig. 2a**). We found that the scores on PC1 were influenced by the nutrient supplements (Kruskal-Wallis test, p-value=0.028), although the scores did not correlate with supplement richness (Spearman’s test, p-value=0.53). This result underscored the complex yet significant impact of nutrient supplements on changes in protein expression levels for most genes. For each gene, plasticity was defined as the standard deviation (SD) of E_pop_ across different nutrient conditions (**Fig. 2b**), representing the mean variability in protein expression levels in response to nutrient changes. We confirmed that plasticity was independent of mean expression levels (Spearman’s test, p-value=0.097) (**Fig. 2c**).

**Fig. 2:**
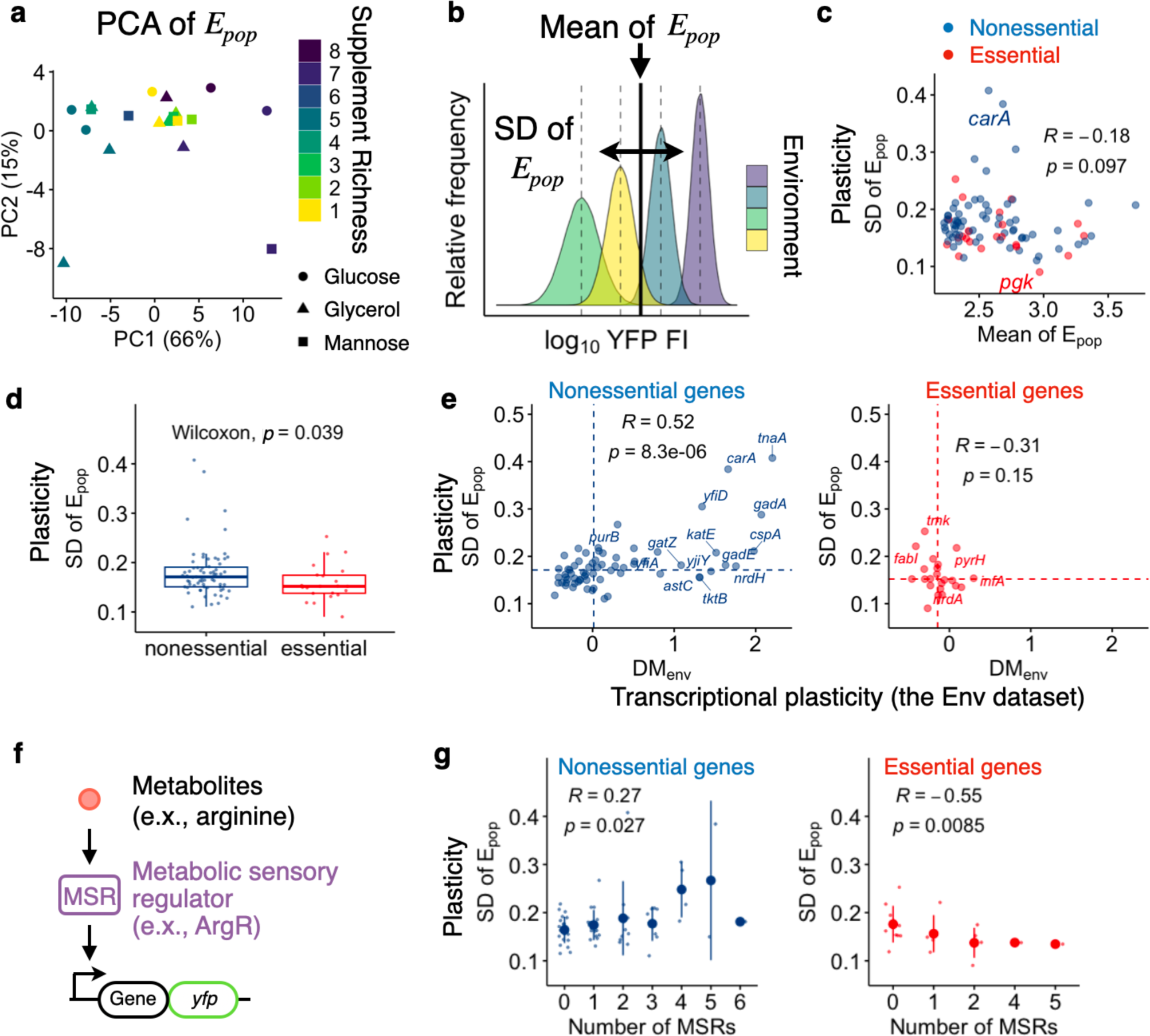
Plasticity in protein expression level across different environments. **a**, First and second principal components (PC) of the mean protein expression levels. Only environmental conditions and genes satisfying defined criteria were subjected to principal component analysis (PCA) as detailed in the **Materials and Methods** section. **b**, Schematic diagram showing the calculation of the mean (vertical solid line) and standard deviation (horizontal arrow, SD) of E_pop_ (vertical dashed lines). The standard deviation of E_pop_ across different environmental conditions was defined as plasticity for each strain. The scale of the horizontal arrow is exaggerated for visualization. **c**, Relationship between the means and standard deviations of E_pop_. Each dot represents a different gene. **d**, Box plots showing the plasticity of essential (red) and nonessential (blue) genes. The inset displays the p-value from the Wilcoxon test. The lower and upper edges of the boxes represent the first (q1) and third (q3) quartiles, respectively. The horizontal lines in the boxes represent the medians (m). The whiskers from the boxes extend to the most extreme observed values inside inner fences, m±1.5(q3–q1). **e**, Relationship between plasticity in mRNA and protein expression levels. Transcriptional plasticity was based on known transcriptome profiles of *E. coli* cultured under different environmental conditions, termed the Env dataset. The DM_env_ was obtained from a previous study and represents the residual standard deviation in mRNA expression levels from the smoothed running median of the standard deviations, as detailed in **Materials and Methods**. **f**, Schematic of regulation by a metabolic sensory regulator (MSR). The activity of MSRs depends on interactions with specific metabolites. **g**, Impact of the number of MSRs on plasticity (left) and noise (right) of the regulated genes. Means and standard deviations are represented by the large points and error bars, respectively. The insets in panels **c**, **e**, and **g** represent Spearman’s R and p-value.

A recent study reported that transcriptional plasticity across different environmental conditions for essential genes tends to be lower than that for nonessential genes^33^. To determine whether a similar trend exists at the level of protein expression, we compared the plasticity between these two categories. We confirmed that essential genes tended to exhibit lower plasticity than nonessential genes (**Fig. 2d**). This common tendency in mRNA and protein expression suggests a contribution of transcriptional plasticity to plasticity at the protein expression levels. To support this, we examined the relationship between transcriptional plasticity, quantified in a previous study as DM_env_, and the SD of E_pop_. The former was based on transcriptome profiles of a single strain of *E. coli* cultured under different environmental conditions (160 profiles, 76 unique environmental conditions), including various nutrient conditions, referred to as the Env dataset. The Env dataset was constructed previously^4^ by removing transcriptome profiles of mutants or other strains from an original large compendium of RNA-seq data in *E. coli*^34,35^. Importantly, all transcriptome profiles were generated within a single laboratory using a standardized protocol, minimizing technical variation within the dataset. We confirmed that our sample set of essential genes exhibited lower transcriptional plasticity than nonessential genes (dashed vertical lines in **Fig. 2e**, Wilcoxon, p-value=0.019), which was consistent with the previous observation for all essential genes in *E. coli*^33^. We found a positive correlation between these two types of plasticity for nonessential genes (Spearman’s rank correlation (R)=0.52, p-value<0.05, **Fig. 2e**), supporting that transcriptional plasticity is retained at the protein expression level for nonessential genes. In contrast, essential genes showed no significant correlation (Spearman, p-value=0.15), likely due to their limited transcriptional plasticity in response to specific environmental changes^33^.

What molecular mechanisms underlie the gene-to-gene difference in plasticity at the protein expression levels? The positive correlation in plasticity between mRNA and protein expression levels suggests that mechanisms responsible for transcriptional plasticity also contribute to plasticity in protein expression levels. A previous study implies that transcriptional regulation by sensory regulators that alter their activity in response to internal and external stimuli, is key to facilitating transcriptional plasticity of their regulatory targets^4^. If this mechanism applies to our conditions, we would expect genes regulated by sensory regulators responsive to changes in nutrient conditions, termed the metabolic sensory regulators (MSR), to exhibit higher plasticity (**Fig. 2f**). To test this hypothesis, we retrieved information about the known relationship between transcriptional regulators (TR) and interacting metabolites from a previous study^36^ (**Supplementary Table S3**), resulting in 69 MSRs with 102 metabolites. We identified 49 of 89 genes of interest regulated by MSR (**Supplementary Table S1**). We found that the plasticity of nonessential genes increased with the number of unique MSRs regulating the genes, supporting our hypothesis (Spearman’s R=0.27, p-value=0.027, **Fig. 2g**). In contrast, the plasticity of essential genes showed the opposite relationship (Spearman’s R=–0.55, p-value=0.0085), suggesting that MSRs are ineffective or even suppressive in facilitating the plasticity of these genes. We also found no significant correlation between plasticity and the number of all transcriptional regulators including MSRs, for nonessential genes (**Supplementary Fig. S1a**), highlighting that MSRs are more critical than general TRs in accounting for the observed plasticity under different nutrient conditions.

### Noise in protein expression levels

Next, we investigated noise in protein expression levels. It is well-stablished that noise depends on the mean expression level^22,25,28^, where noise decreases as mean expression increases at lower levels, mainly due to the dominant contribution of intrinsic noise sources^1,37^. On the other hand, at higher expression levels, noise becomes independent of the mean due to the predominant influence of extrinsic noise sources^1,37^. We confirmed that V_pop_, measured across different nutrient conditions, adhered to these patterns (**Fig. 3a**). To compare noise between genes with different expression levels, it is essential to compensate for the dependency of noise on the mean expression level. Following conventional methods^4,22,25^, we calculated the distance of V_pop_ from a smoothed spline of the running median of V_pop_ (**Fig. 3a**).

**Fig. 3:**
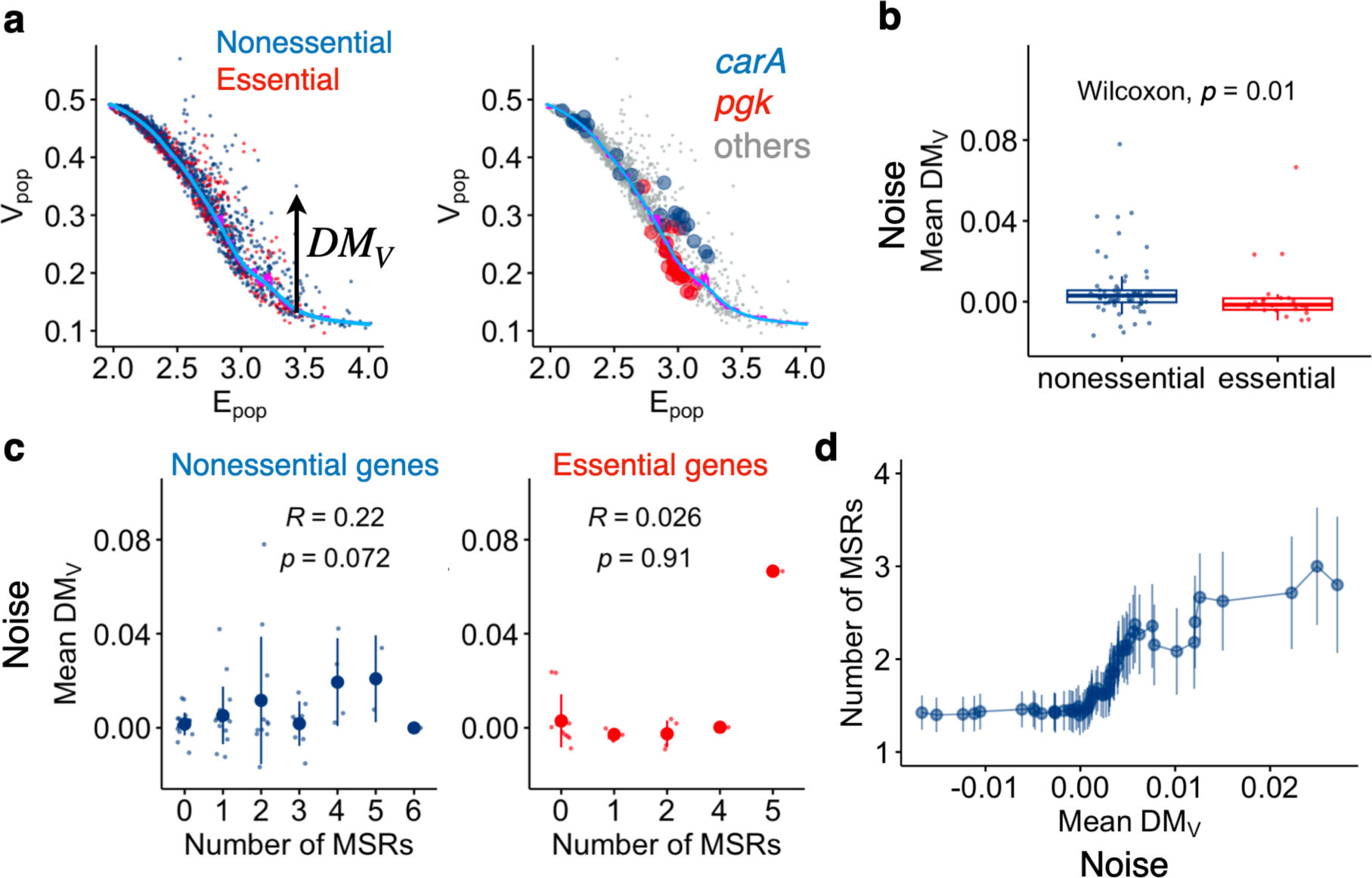
Noise in protein expression level across different environments. **a**, Relationship between V_pop_ and E_pop_ of all genes across all nutrient conditions. Each dot represents a different gene under a specific nutrient condition. The vertical deviation from a smoothed spline (sky blue) calculated from the running median (magenta) of V_p_ is termed DM_V_, representing relative noise in protein expression levels. The right panel highlights two representative genes, *pgk* (red circles) and *carA* (blue circles), from the left panel. Other genes are shown in grey. **b**, Box plots of the mean DM_V_ across different nutrient conditions for essential (red) and nonessential (blue) genes. The lower and upper edges of the boxes represent the first (q1) and third (q3) quartiles, respectively. The horizontal lines in the boxes represent the medians (m). The whiskers from the boxes extend to the most extreme observed values inside inner fences, m±1.5(q3–q1). The inset represents the p-value of the Wilcoxon test. **c**, Relationship between the number of MSRs and noise in protein expression level (mean DM_V_). Means and standard deviations are shown as the large points and the error bars, respectively. Spearman’s R and p-value are shown. **d**, Cumulative analysis of protein noise and MSRs. Genes were ranked according to their noise levels (x) and analyzed in descending order, from high to low noise (left to right on the horizontal axis). For each cut-off on x (horizontal axis), the mean and standard error (vertical axis) of the number of MSRs were calculated for all genes with noise levels exceeding the cut-off value. Due to the inherent nature of cumulative analysis, the initial points (rightmost on the axis) often represent a small number of genes, making both the means and standard errors unreliable. To address this, the first cut-off (rightmost point) was set at the noise level of the sixth gene in the ranked list, ensuring that each data point and corresponding error bar represent more than four genes. Means and standard errors are shown as points and error bars by following Wolf *et al*^11^. This analysis focused on nonessential genes.

A previous study reported that essential genes exhibit lower residual noise compared to the total genes^38^ using the YFP fusion library cultured under a single medium. To determine whether this trend is consistent across different conditions, we computed the mean DM_V_ across various nutrient conditions for each gene (**Supplementary Table S1**) and compared essential and nonessential genes. We confirmed that essential genes exhibited lower DM_V_ than nonessential genes (Wilcoxon test, p-value=0.01) (**Fig. 3b**). Thus, essential genes not only exhibited lower variability in protein expression levels across nutrient conditions but also lower variability in response to stochastic perturbations.

We then examined the impact of the number of MSRs on noise in protein expression levels. A previous study^11^ suggested that promoters with higher noise exhibit larger numbers of regulatory inputs as shown through cumulative analysis. Importantly, however, the study did not conduct a correlation analysis. To explore the relationship between the number of MSRs and noise in protein expression levels, we first performed a standard correlation analysis. Similar to the plasticity in protein expression levels, we found no significant correlation between the number of MSRs and the mean DM_V_ for essential genes (Spearman’s R=0.026, p=0.91, **Fig. 3c right**). For nonessential genes, contrary to the plasticity in protein expression levels, we found no significant positive correlation between the number of MSRs and the mean DM_V_ (Spearman’s R=0.22, p-value=–0.072) (**Fig. 3c left**). Similar results were obtained in the correlation analysis between the mean DM_V_ and the total number of TRs including MSRs (**Supplementary Fig. S1b**). These analyses suggested that the number of MSRs has little detectable impact on noise in protein expression levels for the genes of interest. To improve the sensitivity of detecting a potential weak impact of the number of MSRs on noise, we employed a cumulative method as examined by Wolf *et al*^11^. In this method, genes were first sorted by their noise (DM_V_). For each noise cut-off (x), we calculated the mean and standard error of the number of MSRs for all genes with noise levels above x. By scanning x from high to low noise levels, we generated a cumulative curve representing the relationship between noise and the number of MSRs (**Fig. 3d**). We confirmed a positive association between the number of MSRs and noise in protein expression levels. This analysis also detected the similar associations for all TRs including MSRs (**Supplementary Fig. S2**), consistent with the previous findings^11^. Taken together, our results suggest that genes with higher noise in protein expression levels are associated with a larger number of MSRs, while the contributions to gene-to-gene difference in noise is critically limited.

### Positive correlation between noise and plasticity in protein expression level

Finally, we examined the relationship between plasticity and noise in protein expression levels. Previous studies^23,24^ using datasets of transcriptional plasticity and protein noise have suggested relatively strong positive correlations between these two types of variabilities for nonessential genes (Spearman’s R=0.22 in *E. coli*^23^), while showing no or weaker correlations for essential genes. Despite the interesting implications of these studies, these studies suffered from the discrepancy between mRNA-based plasticity and protein-based noise. To fill the gap, we tested whether the relationships between noise and plasticity, both in protein expression levels, depend on gene essentiality. We found that nonessential genes exhibited a relatively strong positive correlation between noise and plasticity (Spearman’s R=0.61, p-value<0.05), whereas essential genes exhibited no significant correlation (Spearman’s test, p-value=0.25) (**Fig. 4a**). We confirmed these relationships using a conventional method based on transcriptional plasticity (**Fig. 4b**). Thus, noise and plasticity in protein expression levels were positively correlated for nonessential genes, while they appeared incongruent for essential genes.

**Fig. 4:**
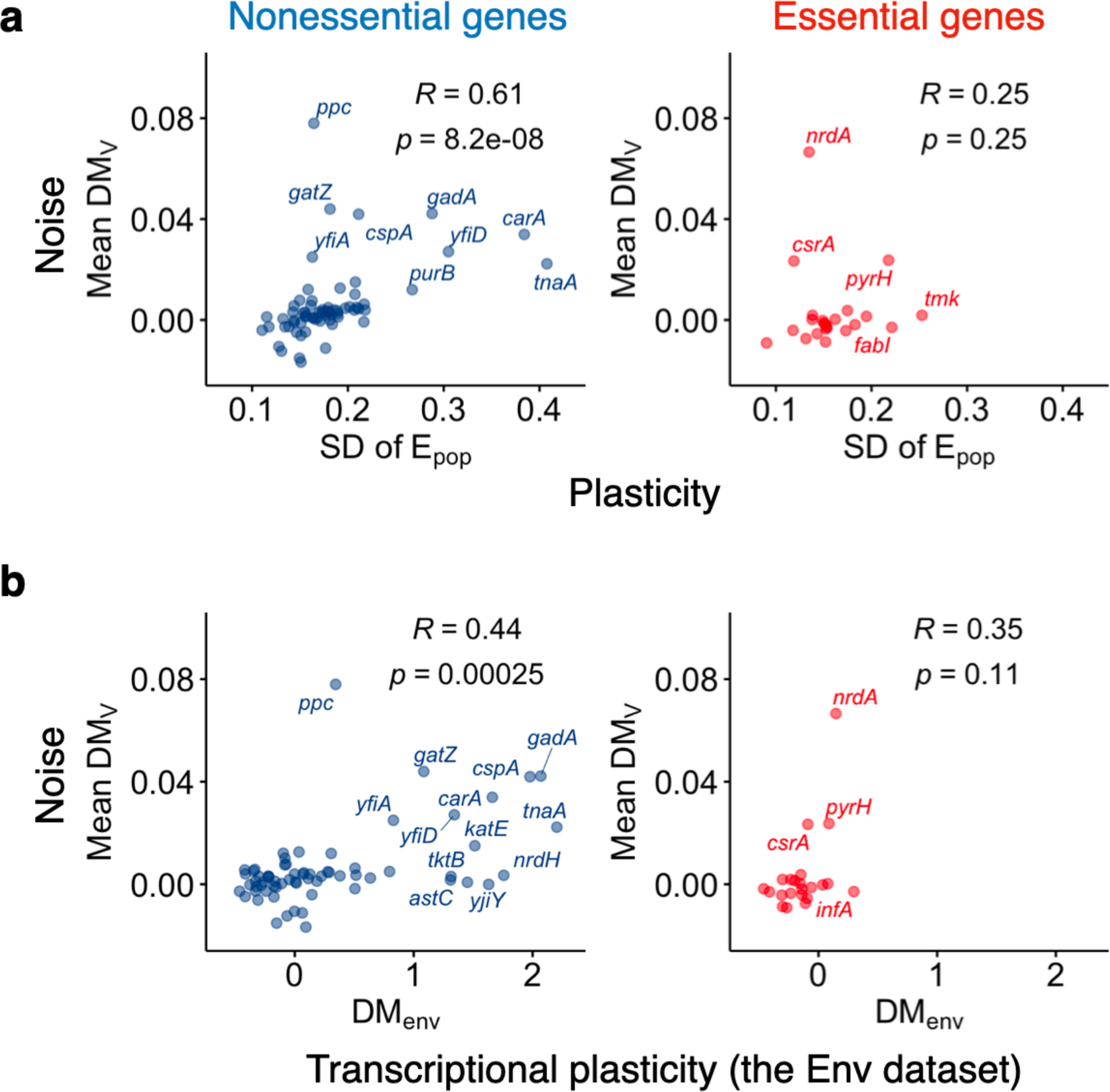
Relationship between noise and plasticity. **a**, Relationship between noise (mean DM_V_) and plasticity (SD of E_pop_) in protein expression levels. **b**, Relationship between noise in protein expression levels and transcriptional plasticity (DM_env_). The left and right panels represent nonessential (left) and essential (right) genes, respectively. The insets represent Spearman’s R and p-value.

## Discussion

Using the YFP fusion library of *E. coli*, we investigated the relationship between noise and plasticity in protein expression levels. Flow cytometry enabled us to measure both the mean and cell-to-cell heterogeneity in protein expression for each gene under different nutrient conditions. We found that noise, plasticity, and their relationship were all dependent on gene essentiality. Essential genes exhibited lower noise and plasticity compared to nonessential genes. Moreover, nonessential genes exhibited a positive correlation between noise and plasticity, a relationship not observed in essential genes. Although this essentiality-dependent relationship had been predicted for several organisms^23,24^, it had not been directly validated at the same gene products until now. Unlike previous methods^11,23^, our novel dataset of plasticity was identical to that of noise, ensuring that the cells shared the same genetic background for these two types of variability. Consequently, our dataset strictly distinguished plasticity from variability in response to genetic mutations, which are often conflated. Additionally, in contrast to previous studies^11,23^, our dataset of protein noise was based on gene expression from natural chromosomal locations in *E. coli*, independent of plasmid copy number variation^39,40^ and potential confounding effects derived from gene expression from plasmids^41^. Despite these stringent considerations relative to the previous study, our results consistently validated the predicted relationships. Thus, this study provides the first direct evidence, using a unified experimental dataset without pooling distinct molecular profiles obtained from separate studies, showing the essentiality-dependent coupling between noise and plasticity in protein expression levels. This experimental validation offers insights into how different types of variability in protein expression levels are organized.

Essential genes, which are crucial for cellular growth, are often under purifying selection or are selected for robustness against different perturbations^42–45^. These selective pressures might contribute to their lower variability in expression levels. What molecular mechanisms could explain the lower plasticity of essential genes? A previous study has shown that essential genes are regulated by a limited number of transcriptional regulators compared to nonessential genes, which may limit their transcriptional variability in response to genetic and environmental perturbations^33^. Consistently, the standard deviation in the number of TRs among essential genes was smaller than that of nonessential genes in our sample sets (**Supplementary Fig. S3b**). This limited number of transcriptional regulators in essential genes might restrict their transcriptional plasticity, thereby limiting their plasticity in protein expression levels (**Fig. 2d**). The scarcity of specific regulatory machineries for reproducible transcriptional responses to perturbations might account for the observed no correlation between noise and plasticity in protein expression levels for essential genes (**Fig. 4**). Another possible mechanism for the lower plasticity of essential genes is that these genes might possess homeostatic transcriptional regulation that helps them withstand metabolic perturbations. In contrast, nonessential genes are reported to have varying numbers of regulators, ranging from none to many transcriptional regulators^33^, a trend confirmed by the standard deviation of the number of TRs among genes in our sample sets (**Supplementary Fig. S3b**). The larger numbers of TRs, particularly MSRs, could contribute to higher transcriptional plasticity in response to metabolic perturbations (**Fig. 2g**), thereby facilitate the plasticity in protein expression levels in these genes. On the other hand, the larger number of regulatory inputs in essential genes might play an opposite role (**Fig. 2g**). Although more regulatory inputs generally lead to higher transcriptional variability due to more target sites^3,4^, some transcriptional regulations can enhance robustness against both genetic and nongenetic perturbations^46,47^. Considering this variation in the roles of transcriptional regulations, the negative correlation between MSR and plasticity in protein expression levels for essential genes suggests that some transcriptional regulations might enhance robustness against metabolic perturbations.

What is the molecular mechanism governing the positive correlation between noise and plasticity in protein expression levels? A global canalization across different perturbations suggests a common molecular mechanism underlying the different types of variability. Typically, a larger number of TRs tends to increase transcriptional variability in response to both genetic and environmental perturbations^3,4^. Consistent with this, genes with a greater number of MSRs tended to exhibit higher plasticity in protein expression (**Fig. 2g**), likely due to a higher probability of sensing nutrient changes in the environment. But how might the number of MSRs contribute to noise in protein expression levels? A previous study proposed that a large number of TRs contribute to facilitate noise in gene expression of their target genes by assuming that the larger the number of TRs, the greater the chance that the target promoter will be sensitive to fluctuations in the activities of at least one of these TRs^48^. This mechanism might partially explain the coupling between noise and plasticity in protein expression levels, considering the associations between noise, plasticity, and the number of MSRs or TRs revealed by a cumulative method (**Fig. 3d**, **Supplementary Fig. S2**). However, our findings suggest that this mechanism alone does not fully account for the positive correlation between noise and plasticity in protein expression levels (**Fig. 3c**, **Supplementary Fig. S1**). Therefore, an alternative common mechanism might explain the congruence between different types of variability, including noise in protein expression levels. Although the current study does not propose a definitive molecular mechanism, there could be a scenario independent of the number of TRs. For example, sensitivity in protein expression of a target gene to fluctuations in TR activity could influence both noise and plasticity in protein expression levels, independent of the number of TRs. Sensitivity, defined as the change in protein expression levels of a regulated gene in response to changes in TR activity, might be a key factor. Higher sensitivities in promoter activity could contribute to greater sensitivity in protein expression levels of downstream genes to fluctuations in TR activity, thereby increasing both noise and plasticity in response to random and nonrandom fluctuations in TRs^49,50^. Other elements involved in gene expression, aside from promoters, might also influence the sensitivity of protein expression levels to both random and nonrandom fluctuations in TR activity. Ideally, future studies should identify the underlying mechanisms connecting plasticity and noise in protein expression levels. Thus, our study provides direct evidence of the expected correlation between noise and plasticity in protein expression levels, while encouraging reconsideration of the dominant mechanism underlying their coupling.

Our study focused on a limited number of genes and environmental conditions. A logical next step is to increase these numbers to explore the generality of our findings across different genes and environmental conditions. Additionally, we addressed only part of the global canalization across different types of variability in protein expression levels. For instance, we did not explore the direct evidence linking noise and mutational variability in protein expression levels. Future studies should address this issue and investigate the molecular mechanisms underlying global canalization across a broader range of variability.

## Materials and Methods

### Bacterial strains and plasmids

We used the YFP fusion library of *E. coli* originally constructed by Taniguchi *et al*^28^. In brief, each strain in the library (totaling 1,023 strains) has a unique YFP-fused ORF on the chromosome. First, we randomly selected 96 strains exhibiting moderate fluorescence levels based on the previous study^28^. The selected strains were transformed with pMW219-CFP, a derivative of the low-copy number pMW219 plasmid carrying cyan fluorescence protein (CFP). The transformants were checked for the absence of localized foci of YFP-fused protein, derived from either aggregation or localization, using microscopy. The CFP fluorescence was used to ensure the cytosol area. The seven strains (*gatY*, *elaB*, *recN*, *aceE*, *yiiU*, *gltD*, and *cheY*) exhibiting localized foci were removed, resulting in 89 strains as listed in **Supplementary Table S1**. These strains and BW25113, used as a control strain without YFP, were subject to flow cytometry. The pMW219-CFP was constructed by assembling pMW219 (NIPPON GENE CO., LTD. Tokyo, Japan, 310-02571) and a CFP expression cassette. The cassette was commercially synthesized (Fasmac Co., Ltd., Japan) and consists of promoter P_LtetO-1_, Shine-Dalgarno sequences SD8, codon-optimized CFP (cerulean) coding sequences, and dual terminators BS2 and T7TE+ as used in Cox *et al*^51^. The integration of the cassette into pMW219 was performed using the In-Fusion HD cloning kit (Takara Bio, Japan, 639648).

### Growth media

We used LB medium (Miller, BD Difco, 244620) for the construction of bacterial cells and precultures for flow cytometry. We used a synthetic medium, a derivative of M9 minimal medium (pH 7.2), as a basal medium containing 47.7 mM Na_2_HPO_4_ (Disodium Hydrogenphosphate 12-Water, Wako, Japan, 196-02835), 22.0 mM KH_2_PO_4_ (Potassium Dihydrogenphosphate, Wako, Japan, 164-22635), 8.6 mM NaCl (Sodium Chloride, Wako, Japan, 191-01665), 18.7 mM NH_4_Cl (Ammonium Chloride, Wako, Japan, 014-03005), 2 mM MgSO_4_ (Magnesium Sulfate Heptahydrate, Wako, Japan, 138-00415), 0.1 mM CaCl_2_ (Calcium Chloride Dihydrate, Wako, Japan, 038-24985), 10 mM FeSO_4_ (Iron(II) Sulfate Heptahydrate, Wako, Japan, 094-01082) and 4 µM biotin ((+)-Biotin, Wako, Japan, 023-08711), with supplementation of 25 µg/mL Kanamycin (Kanamycin Sulfate, Wako, Japan, 115-00342) when required. We used glucose (D(+)-Glucose, Wako, Japan, 049-31165), glycerol (Glycerol, Wako, Japan, 075-00616), and mannose (D(+)-Mannose, Wako, Japan, 132-00871) as single carbon sources. Additionally, we constructed eight categories of supplements: casamino acids, 20 amino acids, 12 amino acids, two cocktails of six amino acids (6A, 6B), nucleotides, vitamins, and trace elements as detailed in **Supplementary Table S2**.

### Culture conditions

Cells were inoculated from frozen stocks into 200 μL/well of LB medium in 96-well microplates (Greiner Bio-One, Cellstar, 655180). The bacterial cultures were incubated at 32 °C with shaking at 800 rpm overnight using an incubator (TAITEC, Saitama, Japan, MBR-034P). The grown cultures were diluted 100-fold with M9 buffer and 2 μL of the dilutions were transferred into 200 μL/well of fresh synthetic medium in 96-well microplates with a 96-channel pipette (PLATEMASTER, Gilson, USA). The bacterial cultures were incubated at 32 °C with shaking at 800 rpm overnight. The cell density of the grown cultures was measured using a plate reader (Tecan, Switzerland, Infinite F200). The bacterial cultures were diluted 100-fold with M9 buffer and transferred into 200 μL/well of fresh synthetic medium in 96 well microplates. The amount of inoculum was determined so that the subsequent overnight cultures remained in the exponential phase, using the growth characteristics of the corresponding conditions obtained preliminarily. The bacterial cultures were incubated at 32 °C with shaking at 800 rpm overnight. The cell density of the overnight cultures was measured to ensure the designed growth phases. The bacterial cultures were diluted with M9 buffer to reach an appropriate cell density for flow cytometry. Two biological replicates were cultured on different days.

### Flow cytometry

Cells dispensed in M9 buffer were analyzed using a flow cytometer (BD, USA, FACSAria III). Fluorescence intensity from YFP (YFP FI) was measured using a 488-nm argon laser and a 515–545 nm emission filter. The following PMT voltage settings were used: forward scatter (FSC), 350; side scatter (SSC), 350; yellow fluorescence: 800. The events exhibiting higher values in FSC and SSC than the thresholds (200 for each) were recorded. A total of 10,000–50,000 events were recorded for each population.

### Data analysis in flow cytometry

The narrow gates of FSC were determined for each environmental condition each measurement date. First, all FCS files obtained under the same environmental conditions on the same dates were read using the flowCore^52^ package in R^53^ to construct cumulative populations. The modes of log_10_-transformed FSC of the cumulative populations were calculated subsequently. The narrow FSC gates were defined as the range between the modes ± 0.1 on the logarithmic scale. The events within the gates were subjected to subsequent analysis. The autofluorescence intensity derived from BW25113 without YFP was subtracted from YFP FI. The events exhibiting lower values in raw YFP FI below the autofluorescence intensity were removed. Populations with fewer than 4,000 events after the above filtering were omitted from the subsequent analysis. Environmental conditions lacking biological duplicates were also omitted. The means (E_pop_) and standard deviations (V_pop_) in log_10_-transformed YFP FI were calculated for each population. These statistics were averaged among the biological replicates. Plasticity in protein expression levels for individual genes was defined as the standard deviations of E_pop_ of the corresponding strains among the different nutrient conditions. To compensate for the E_pop_-dependency of V_pop_, we calculated the distance of each standard deviation from a smoothed running median of standard deviations, referred to as DM_V_, following the method outlined in previous studies^25,54^. The means of DM_V_ were calculated among the different nutrient conditions.

### Compilation of the regulatory interactions between genes

The interactions between metabolites and transcription factors were obtained from a previous study^36^. The transcription factors interacting with metabolites were termed metabolic sensory regulators (MSR, **Supplementary Table S3**). The known regulatory interactions between 69 MSRs and the regulatory target genes were obtained from RegulonDB 11.0^55^ and EcoCyc^56^.

### Principal component analysis

Some nutrient conditions downregulated many genes below the detection limit in flow cytometry, which greatly reduced the number of genes applicable to PCA. To avoid this situation, nutrient conditions that reduced more than four genes applicable to PCA were excluded first. Subsequently, the genes that were applicable to PCA for all the filtered conditions were used. As a result, 69 genes with 19 nutrient conditions were subjected to PCA. PCA was conducted using the prcomp function in R.

### Cumulative analysis for the relationship between noise, plasticity, and the number of transcriptional regulators

Nonessential genes were ranked according to their noise levels (mean DM_V_) and analyzed in a descending order. As a function of a cut-off on noise (x), the mean and standard error of the number of MSRs (or TRs) were calculated for all nonessential genes with noise levels above the cut-off value. A similar approach was applied for plasticity, where nonessential genes were again ranked according to their noise levels and scanned from high to low. For each cut-off on noise (x), the mean and standard error of plasticity (mean E_pop_) were calculated for all nonessential genes with noise levels exceeding the cut-off value.

### Data visualization

All figures were generated in R. Illustrative plots were created using the ggplot2^57^ and ggpubr^58^ packages.

## Data availability

The raw fcs fata are available at Zenodo (https://doi.org/10.5281/zenodo.13318136; https://doi.org/10.5281/zenodo.13318501; https://doi.org/10.5281/zenodo.13319277).

## Code availability

Custom R scripts used in the manuscript are available at GitHub (https://github.com/tsuruubi/Noise_Plasticity_E_Coli).

## Ethics declarations

The authors declare no competing interests.

## Acknowledgements

The authors thank H. Koike for technical supports in flow cytometry. This work was supported by, the Japan Society for Promotion of Science (JSPS) KAKENHI (18H02427, 22H05403, 24K21985 to S.T.;17H06389 to C.F. and S.T.; 22K21344 to C.F.), Japan Science and Technology Agency (JST) ERATO (JPMJER1902 to S.T. and C.F.).

## References

1. Elowitz, M.B., Levine, A.J., Siggia, E.D. & Swain, P.S. Stochastic gene expression in a single cell. Science 297, 1183–6 (2002).

2. Tirosh, I., Weinberger, A., Carmi, M. & Barkai, N. A genetic signature of interspecies variations in gene expression. Nat Genet 38, 830–4 (2006).

3. Landry, C.R., Lemos, B., Rifkin, S.A., Dickinson, W.J. & Hartl, D.L. Genetic properties influencing the evolvability of gene expression. Science 317, 118–21 (2007).

4. Tsuru, S. & Furusawa, C. Genetic properties underlying transcriptional variability across different perturbations. bioRxiv, 2024.04.15.589659 (2024).

5. Denver, D.R. et al. The transcriptional consequences of mutation and natural selection in *Caenorhabditis elegans*. Nat Genet 37, 544–8 (2005).

6. Uller, T., Moczek, A.P., Watson, R.A., Brakefield, P.M. & Laland, K.N. Developmental Bias and Evolution: A Regulatory Network Perspective. Genetics 209, 949–966 (2018).

7. Milocco, L. & Uller, T. Utilizing developmental dynamics for evolutionary prediction and control. Proc Natl Acad Sci U S A 121, e2320413121 (2024).

8. Brun-Usan, M., Rago, A., Thies, C., Uller, T. & Watson, R.A. Development and selective grain make plasticity ‘take the lead’ in adaptive evolution. BMC Ecol Evol 21, 205 (2021).

9. Brun-Usan, M., Zimm, R. & Uller, T. Beyond genotype-phenotype maps: Toward a phenotype-centered perspective on evolution. Bioessays 44, e2100225 (2022).

10. Levis, N.A. & Pfennig, D.W. Evaluating ‘Plasticity-First’ Evolution in Nature: Key Criteria and Empirical Approaches. Trends Ecol Evol 31, 563–574 (2016).

11. Wolf, L., Silander, O.K. & van Nimwegen, E. Expression noise facilitates the evolution of gene regulation. Elife 4(2015).

12. Eldar, A. et al. Partial penetrance facilitates developmental evolution in bacteria. Nature 460, 510–4 (2009).

13. Meiklejohn, C.D. & Hartl, D.L. A single mode of canalization. Trends in Ecology & Evolution 17, 468–473 (2002).

14. Kaneko, K. Relationship among phenotypic plasticity, phenotypic fluctuations, robustness, and evolvability; Waddington’s legacy revisited under the spirit of Einstein. J Biosci 34, 529–42 (2009).

15. Furusawa, C. & Kaneko, K. Formation of dominant mode by evolution in biological systems. Phys Rev E 97, 042410 (2018).

16. Draghi, J.A. & Whitlock, M.C. Phenotypic plasticity facilitates mutational variance, genetic variance, and evolvability along the major axis of environmental variation. Evolution 66, 2891–902 (2012).

17. Waddington, C.H. Genetic assimilation of an acquired character. Evolution, 118–126 (1953).

18. Wagner, G.P., Booth, G. & Bagheri-Chaichian, H. A Population Genetic Theory of Canalization. Evolution 51, 329–347 (1997).

19. Rifkin, S.A., Houle, D., Kim, J. & White, K.P. A mutation accumulation assay reveals a broad capacity for rapid evolution of gene expression. Nature 438, 220–3 (2005).

20. Noble, D.W.A., Radersma, R. & Uller, T. Plastic responses to novel environments are biased towards phenotype dimensions with high additive genetic variation. Proc Natl Acad Sci U S A 116, 13452–13461 (2019).

21. Cheverud, J.M. A Comparison of Genetic and Phenotypic Correlations. Evolution 42, 958–968 (1988).

22. Silander, O.K. et al. A genome-wide analysis of promoter-mediated phenotypic noise in *Escherichia coli*. PLoS Genet 8, e1002443 (2012).

23. Singh, G.P. Coupling between noise and plasticity in *E. coli*. G3 (Bethesda) 3, 2115–20 (2013).

24. Lehner, B. Conflict between noise and plasticity in yeast. PLoS Genet 6, e1001185 (2010).

25. Newman, J.R. et al. Single-cell proteomic analysis of *S. cerevisiae* reveals the architecture of biological noise. Nature 441, 840–6 (2006).

26. Mustoe, A.M. et al. Pervasive Regulatory Functions of mRNA Structure Revealed by High-Resolution SHAPE Probing. Cell 173, 181–195 e18 (2018).

27. Guimaraes, J.C., Rocha, M. & Arkin, A.P. Transcript level and sequence determinants of protein abundance and noise in *Escherichia coli*. Nucleic Acids Res 42, 4791–9 (2014).

28. Taniguchi, Y. et al. Quantifying *E. coli* proteome and transcriptome with single-molecule sensitivity in single cells. Science 329, 533–8 (2010).

29. Si, F. et al. Invariance of Initiation Mass and Predictability of Cell Size in *Escherichia coli*. Curr Biol 27, 1278–1287 (2017).

30. Goodall, E.C.A. et al. The essential genome of *Escherichia coli* K-12. mBio 9, 1–18 (2018).

31. Tsuru, S. et al. Noisy cell growth rate leads to fluctuating protein concentration in bacteria. Phys Biol 6, 036015 (2009).

32. Tsuru, S. et al. Adaptation by stochastic switching of a monostable genetic circuit in Escherichia coli. Mol Syst Biol 7, 493 (2011).

33. Tsuru, S., Hatanaka, N. & Furusawa, C. Promoters constrain evolution of expression levels of essential genes in *Escherichia coli*. bioRxiv, 2024.05.20.594948 (2024).

34. Sastry, A.V. et al. The *Escherichia coli* transcriptome mostly consists of independently regulated modules. Nat Commun 10, 5536 (2019).

35. Rychel, K. et al. iModulonDB: a knowledgebase of microbial transcriptional regulation derived from machine learning. Nucleic Acids Res 49, D112–D120 (2021).

36. Lempp, M. et al. Systematic identification of metabolites controlling gene expression in *E. coli*. Nat Commun 10, 4463 (2019).

37. Swain, P.S., Elowitz, M.B. & Siggia, E.D. Intrinsic and extrinsic contributions to stochasticity in gene expression. Proc Natl Acad Sci U S A 99, 12795–800 (2002).

38. Fuentes, D.A.F., Manfredi, P., Jenal, U. & Zampieri, M. Pareto optimality between growth-rate and lag-time couples metabolic noise to phenotypic heterogeneity in *Escherichia coli*. Nat Commun 12, 3204 (2021).

39. Shao, B. et al. Single-cell measurement of plasmid copy number and promoter activity. Nat Commun 12, 1475 (2021).

40. Paulsson, J. Summing up the noise in gene networks. Nature 427, 415–8 (2004).

41. Brewster, R.C. et al. The transcription factor titration effect dictates level of gene expression. Cell 156, 1312–1323 (2014).

42. Ho, W.C. & Zhang, J. Adaptive Genetic Robustness of *Escherichia coli* Metabolic Fluxes. Mol Biol Evol 33, 1164–76 (2016).

43. Cohen, O., Oberhardt, M., Yizhak, K. & Ruppin, E. Essential Genes Embody Increased Mutational Robustness to Compensate for the Lack of Backup Genetic Redundancy. PLoS One 11, e0168444 (2016).

44. Shibai, A., Kotani, H., Sakata, N., Furusawa, C. & Tsuru, S. Purifying selection enduringly acts on the sequence evolution of highly expressed proteins in *Escherichia coli*. G3 (Bethesda) 12(2022).

45. Hawkins, J.S. et al. Mismatch-CRISPRi Reveals the Co-varying Expression-Fitness Relationships of Essential Genes in *Escherichia coli* and *Bacillus subtilis*. Cell Syst 11, 523–535 e9 (2020).

46. Denby, C.M., Im, J.H., Yu, R.C., Pesce, C.G. & Brem, R.B. Negative feedback confers mutational robustness in yeast transcription factor regulation. Proc Natl Acad Sci U S A 109, 3874–8 (2012).

47. Becskei, A. & Serrano, L. Engineering stability in gene networks by autoregulation. Nature 405, 590–3 (2000).

48. Urchueguia, A. et al. Genome-wide gene expression noise in Escherichia coli is condition-dependent and determined by propagation of noise through the regulatory network. PLoS Biol 19, e3001491 (2021).

49. Hooshangi, S., Thiberge, S. & Weiss, R. Ultrasensitivity and noise propagation in a synthetic transcriptional cascade. Proc Natl Acad Sci U S A 102, 3581–6 (2005).

50. Becskei, A., Kaufmann, B.B. & van Oudenaarden, A. Contributions of low molecule number and chromosomal positioning to stochastic gene expression. Nat Genet 37, 937–44 (2005).

51. Cox, R.S., 3rd, Dunlop, M.J. & Elowitz, M.B. A synthetic three-color scaffold for monitoring genetic regulation and noise. J Biol Eng 4, 10 (2010).

52. Ellis, B., et al. flowCore: flowCore: Basic structures for flow cytometry data. R package version 2.14.2, https://bioconductor.org/packages/flowCore/ (2024).

53. R Core Team. R: A Language and Environment for Statistical Computing. (2023).

54. Uchida, Y., Shigenobu, S., Takeda, H., Furusawa, C. & Irie, N. Potential contribution of intrinsic developmental stability toward body plan conservation. BMC Biol 20, 82 (2022).

55. Tierrafria, V.H. et al. RegulonDB 11.0: Comprehensive high-throughput datasets on transcriptional regulation in *Escherichia coli* K-12. Microb Genom 8(2022).

56. Keseler, I.M. et al. The EcoCyc Database in 2021. Front Microbiol 12, 711077 (2021).

57. Wickham, H. ggplot2: Elegant Graphics for Data Analysis. https://ggplot2.tidyverse.org (2016).

58. Kassambara, A. ggpubr: ‘ggplot2’ Based Publication Ready Plots. R package version 0.6.0, https://rpkgs.datanovia.com/ggpubr/ (2023).

