## Supplementary Fig for "Congruence between noise and plasticity in protein expression"

**I. Supplementary Figures**
**II. Supplementary References**

### I. Supplementary Figures

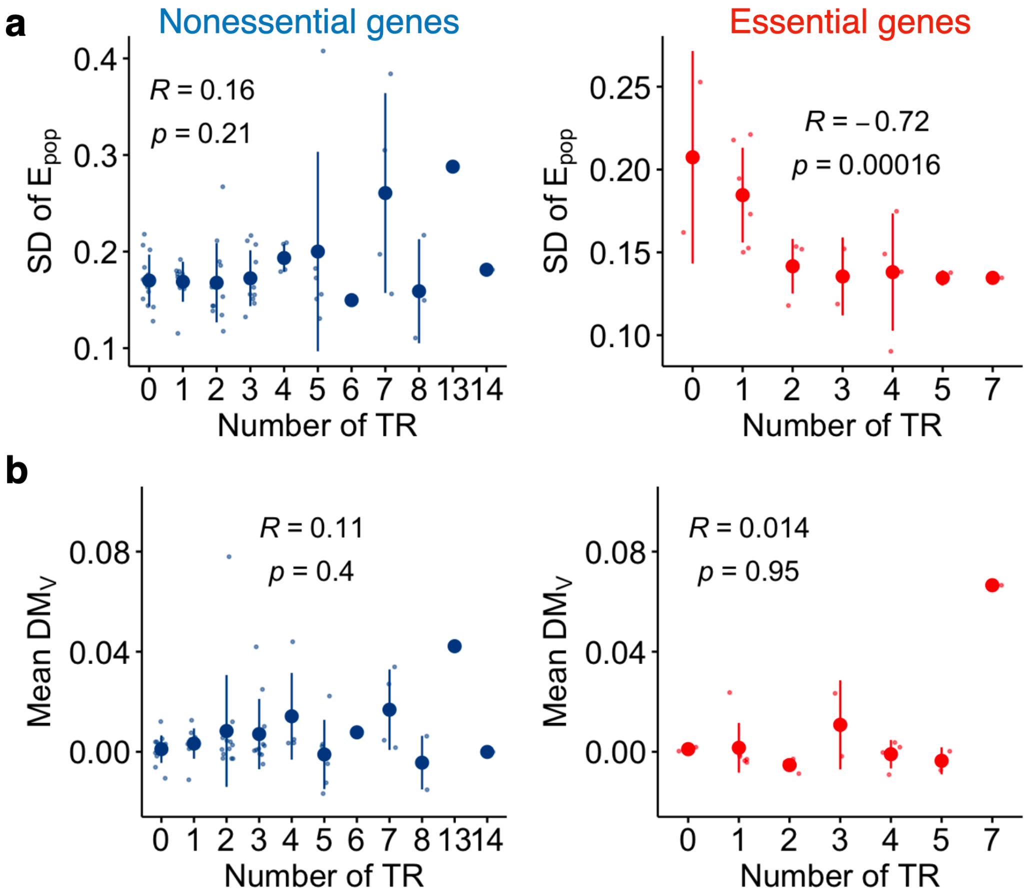


**Fig. S1: Relationship between the number of transcriptional regulators, plasticity, and noise**

**a**, Relationship between the number of transcriptional regulators (TR) and plasticity in protein expression levels (SD of $E_{pop}$). All transcriptional regulators including MSRs and other TRs were considered. **b**, Relationship between the number of TRs and noise in protein expression level (mean $DM_V$). Means and standard deviations are shown as the large points and the error bars, respectively. Spearman's R and p-value are shown.

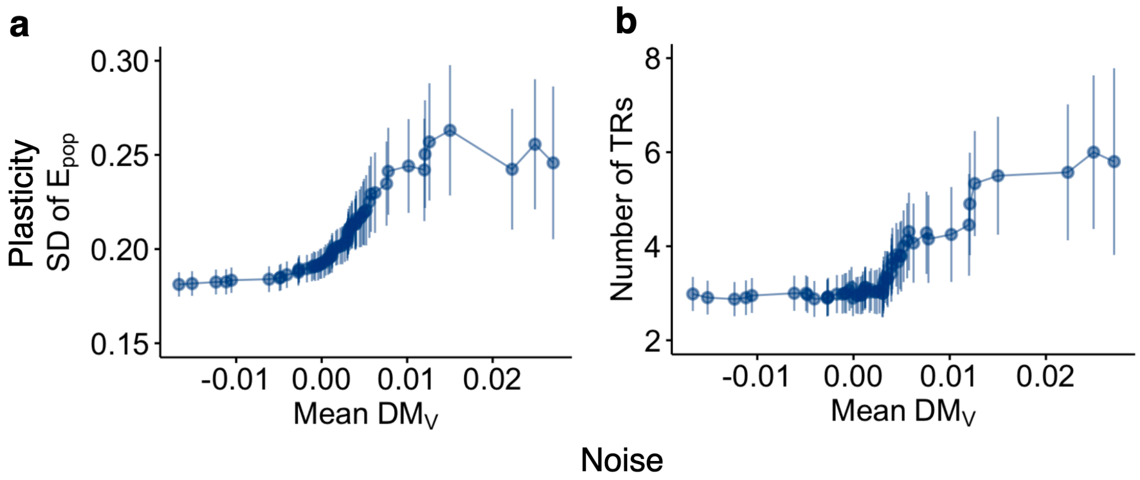


**Fig. S2: A cumulative analysis for relationship between the number of transcriptional regulators, plasticity, and noise**

Following a previous study (Figure 2 in Wolf *et al*[1]), a cumulative approach was employed to explore the relationship between the number of transcriptional regulators, plasticity, and noise in protein expression level. **a**, Genes were sorted by their noise level (x) As a function of a cut-off on x (horizontal axis), the mean and standard error (vertical axis) of plasticity in protein expression levels were calculated for all genes with the noise greater than x. **b**, Genes were sorted by their noise level x as in panel **a**, and the mean and standard error of the number of TRs including MSRs were calculated for all genes with noise level greater than x. The first cut-off (rightmost point) was set at the noise level of the sixth gene in the sorted list to ensure that each data point and error bar correspond to more than four genes for all panels. Means and standard errors are shown as the points and error bars. Nonessential genes were considered.

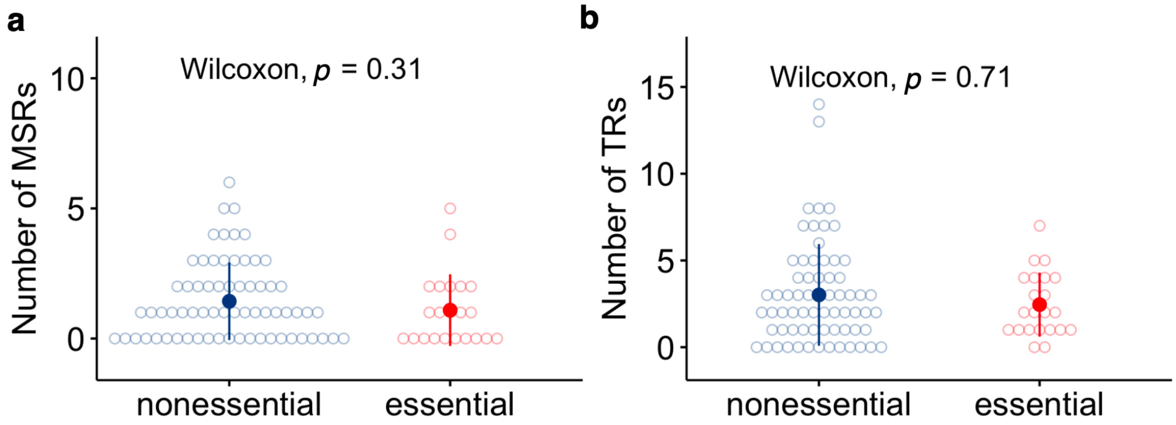


**Fig. S3: Relationship between the number of transcriptional regulators and essentiality**

**a**, Relationship between the number of MSRs and essentiality of target genes for cellular growth. **b**, Relationship between the number of all TRs, including MSRs, and essentiality of target genes. Means and standard deviations are shown as the large points and the error bars, respectively.

#### II. Supplementary References

1. Wolf, L., Silander, O.K. & van Nimwegen, E. Expression noise facilitates the evolution of gene regulation. *Elife* **4**(2015).
